# Single-molecule sequencing of animal mitochondrial genomes reveals chloroplast-like architecture and repeat-mediated recombination

**DOI:** 10.1101/2022.08.11.503648

**Authors:** Joel Sharbrough, Laura Bankers, Emily Cook, Peter D. Fields, Joseph Jalinsky, Kyle E. McElroy, Maurine Neiman, John M. Logsdon, Jeffrey L. Boore

**Affiliations:** Department of Biology, New Mexico Institute of Mining and Technology, Socorro, NM 87801; Department of Biology, University of Iowa, Iowa City, IA 52242, USA; Zoologisches Institut, University of Basel, Vesalgasse 1, 4051 Basel, Switzerland; Department of Ecology, Evolution, and Organismal Biology, Iowa State University, IA 50011, USA; Phenome Health and Institute for Systems Biology, 401 Terry Avenue N, Seattle, WA 98109, USA

**Keywords:** Flip-flop recombination, heteroplasmy, mitochondrial DNA, PacBio sequencing, repeat-mediated recombination

## Abstract

Recent advances in long-read sequencing technology have allowed for single-molecule sequencing of entire mitochondrial genomes, opening the door for direct investigation of mitochondrial genome architecture and landscapes of recombination. We used PacBio sequencing to re-assemble mitochondrial genomes from two species of New Zealand freshwater snails, *Potamopyrgus antipodarum* and *Potamopyrgus estuarinus*. These assemblies revealed a ∼1.7 kb structure within the mitochondrial genomes of both species that was previously undetected by assembly of short sequencing reads and likely corresponding to a large non-coding region commonly present in mitochondrial genomes. The overall architecture of these *Potamopyrgus* mitochondrial genomes is reminiscent of the chloroplast genomes of land plants, harboring a large single-copy region (LSC) and a small single-copy region (SSC) separated by a pair of inverted repeats (IRa and IRb). Individual sequencing reads that spanned across the *Potamopyrgus* IRa–SSC–IRb structure revealed the occurrence of “flip-flop” recombination, apparently mediated by the IRs. We also detected evidence for two distinct IR haplotypes and recombination between them in wild-caught *P. estuarinus*, as well as extensive inter-molecular recombination between SNPs in the LSC region. Together, these observations suggest that mitochondrial inheritance is not strictly maternal in these snails. The chloroplast-like architecture and repeat-mediated mitochondrial recombination we describe here raise fundamental questions regarding the origins and commonness of such architecture, whether and how recombination mediates mitochondrial genome evolution, and the role of genome architecture in driving cytoplasmic genome biology and the maintenance of cytoplasmic genomes.

## Introduction

Wide taxonomic sampling enabled by the genomic era has revealed remarkable diversity in the content, architecture, and biology of mitochondrial genomes (1–3). The mitochondrial genomes of bilaterian animals have been particularly well studied. With few (but notable) exceptions, these genomes are of small size (∼16-17 kb) (4), encode only a tiny fraction of the genes necessary to carry out tasks performed within mitochondria (4), engage in complex co-evolution with their nuclear-encoded interacting partners (5–8), have relatively high mutation rates (9, 10) and seemingly reduced levels of homologous recombination (11), and are predominantly maternally inherited (12, 13).

Bilaterian animal mitochondrial DNAs (mtDNA) are typically very compact, with few intergenic nucleotides except for a single non-coding region ranging in size from several dozen to several thousand nucleotides (14). In some cases, this non-coding region also contains a third DNA strand that causes a displacement of base pairing in this region, the so-called “D-loop” structure (15). Hereafter we will refer to this non-coding region as the “D-loop region” regardless of whether a D-loop structure has been observed for any particular taxon. The D-loop region has been shown to contain regulatory elements for initiation, termination of transcription, and replication in several taxa (16). This structure and function are presumed to exist across animals, although obvious similarities in primary sequence or potential secondary structures are not readily identifiable across distantly related taxa. This apparent lack of homology is probably related to the fact that portions of the D-loop region are often hypervariable and can contain repeated sequence elements (17–20).

The variability and repetitive nature of the D-loop region likely also contribute to why it has been especially difficult to amplify by PCR in many animal whole-mitochondrial genome sequencing efforts. Difficulties in PCR amplification of this region could be due to its highly biased base composition, the presence of secondary structures that are difficult for the polymerase to read through, or the response of the polymerase, itself a prokaryotic DNA polymerase, to signals for terminating replication of the mtDNA. Barriers to sequencing posed by the D-loop region could also explain why many animal mitochondrial genomes, even though listed as “complete” and “circular” in GenBank, do not report the D-loop region sequence (*e.g.*, (21–28)).

Almost all mitochondrial genome assemblies have relied on bioinformatically assembling sequencing reads generated *via* Sanger sequencing (∼700 nucleotides (nt) in length) or next-generation sequencing (NGS; 100-250 nt in length) from preparations of whole DNA or that are enriched for mtDNA. This strategy often fails to successfully assemble repeated elements of a length larger than any individual read that cannot find a unique flanking sequence on each side. This problem is exacerbated by assembly *via* de Bruijn graphs, which provide computationally tractable assembly of millions of short reads (*e.g*., (29–33)), but also typically break assemblies at complex structural features (34) such as those common in mitochondrial D-loop regions.

The so-called “third-generation sequencing” reads (*i.e*., PacBio and Oxford Nanopore) that can span many kilobases offer the tantalizing opportunity to uncover previously undetectable genomic architectural features. The application of these long-read technologies has already led to major advances in the identification of disease-causing variants (35), the generation of complete microbial genome assemblies (36), the detection of rare mutations and DNA damage (37), the production of assembly-free transcriptomes (38), the description of the complex patterns of mitochondrial recombination in plant mitochondria (39, 40), and a truly complete human genome sequence with each chromosome in a single, uninterrupted contig from one telomere to the other (41).

The New Zealand freshwater snail, *Potamopyrgus antipodarum* (Figure 1), is a powerful model for a number of ecological and evolutionary questions, including the maintenance of sex (42), consequences of polyploidy (43), host-parasite dynamics (44–46), biology of invasive organisms (47), and mitonuclear coevolution (25, 28, 48, 49). To generate the whole-genome resources needed to investigate these fundamental topics, we used a combination of Illumina HiSeq (DNA and RNA), MiSeq, and PacBio long-read technologies to sequence total cellular DNA from an inbred (∼25 generations) sexual lineage of *P. antipodarum* and wild-caught specimens of its close relative ((50), our data), *P. estuarinus*. Here, we used a subset of these sequencing reads to re-assemble the mitochondrial genomes of both species. Our complete and circular mitochondrial assemblies report a novel ∼1.7 kb repeat structure not detected in previous whole-mitochondrial genome sequencing efforts (25, 28) associated with intra- and inter-molecular recombination. Long PacBio reads from *P. antipodarum* were able to detect two distinct orientations of the 1.7kb repeat at relatively high abundance, reminiscent of a similar observation first made in chloroplast genomes (51), potentially implicating flip-flop recombination as a mechanism of intra-molecular mitochondrial recombination. We also observed high levels of SNP diversity in wild-caught *P. estuarinus* samples and numerous recombinant molecules in pooled-MiSeq data, implying heteroplasmy occurs with high frequency in the population. Indeed, the abundance of recombinant molecules in such a small sample of individuals leads us to speculate that *Potamopyrgus* exhibits high degrees of paternal leakage or even biparental inheritance of mitochondrial DNA. Together, these observations provide critical steps towards understanding the biology of this fascinating natural system, with potentially broad implications for mitochondrial biology, inheritance, and evolution in animals.

**Figure 1.**
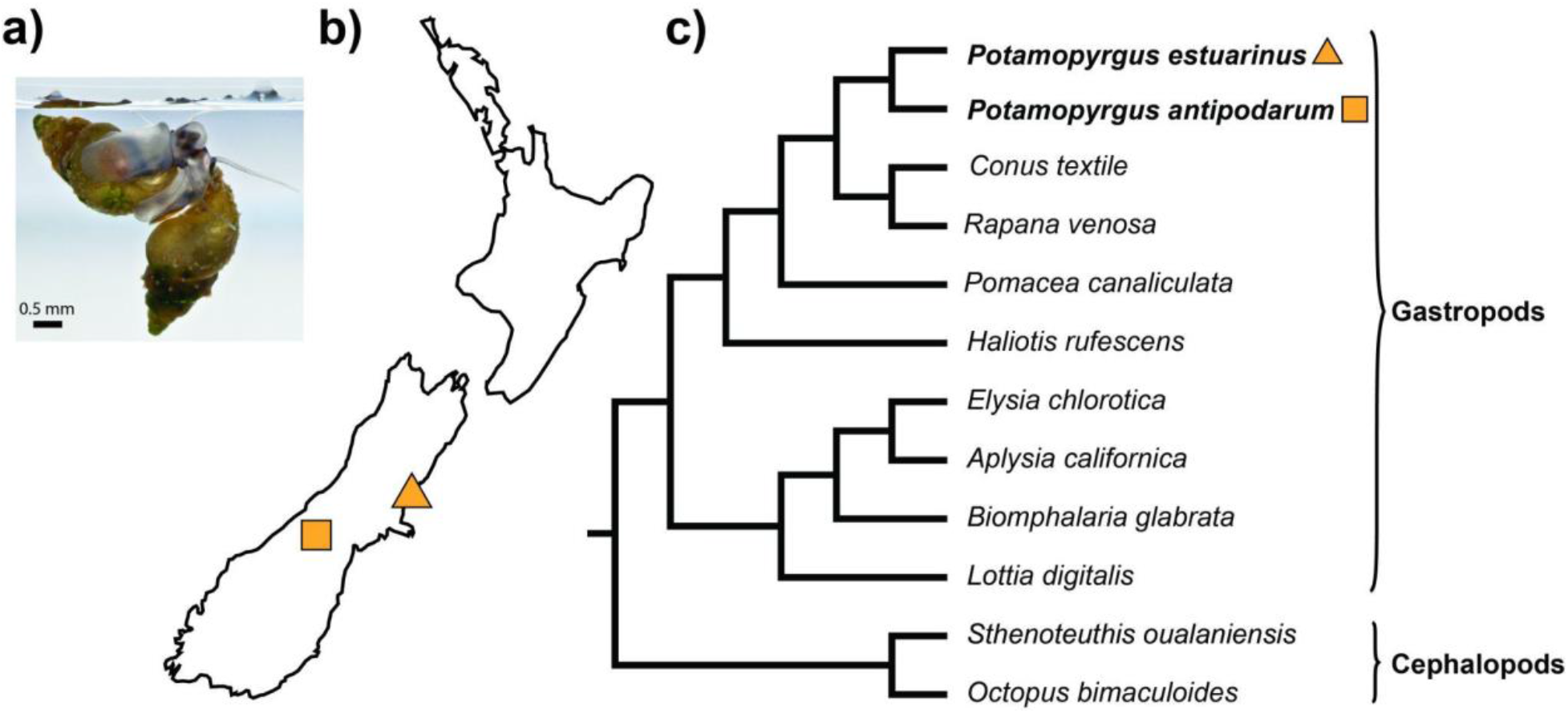
*Potamopyrgus antipodarum* and *P. estuarinus* (*Caenogastropoda*) are snails native to New Zealand. a) *Potamopyrgus antipodarum* is approximately 4-6 mm in length. b) Map of New Zealand depicting sampling locations of the snails used in this study, and a scaled image of *P. estuarinus*. b) Cladogram of selected mollusc species depicting *Potamopyrgus* within the *Caenogastropoda*.

## Results and Discussion

### Mitochondrial genome architecture in Potamopyrgus resembles chloroplast genomes

Despite very high coverage (>800x, see below), our first efforts to assemble complete and circular *Potamopyrgus* mitochondrial genomes using Illumina HiSeq paired-end reads with SPAdes v. 3.13.0 (31) were unsuccessful, with the mitochondrial contigs from the de Bruijn assembly graphs forming “barbell” (*P. antipodarum*) and “lollipop” (*P. estuarinus*) structures (Figure S1) rather than simple circles. Additional attempts to polish and circularize the assembly with Pilon v 1.23 (52) were unsuccessful, but we confirmed that the region preventing circularization corresponded exactly to the barbell and lollipop structural anomalies. Using PacBio reads from both species (acquired as part of the ongoing *P. antipodarum* and *P. estuarinus* nuclear genome sequencing projects), we discovered a previously undetected structural element of ∼1.7 kb in length. In both species, the structure consists of a pair of inverted repeats (IRs) interrupted by a small single-copy region (SSC). In the mtDNAs of *P. antipodarum*, IRa and IRb are identical in sequence, and the intervening SSC consists of two dinucleotide repeats (TA[x54] and TC[x337]). By contrast, we detected 15 high-confidence SNPs (*i.e.*, present in both PacBio and MiSeq pool-seq datasets) in the IRs of wild-caught *P. estaurinus*, all of which can be observed in both IRs, and the intervening SSC consists solely of a TA[x334] dinucleotide repeat.

Because the TA[x334] dinucleotide repeat that forms the entirety of the SSC in *P. estuarinus* is a palindrome, its orientation relative to the LSC is impossible to determine. However, the *P. antipodarum* SSC does have identifiable directionality because it is made up of a series of two dinucleotide repeats (TA[x54] and TC[x337]) that, when reversed, read GA[x337] and TA[x54], respectively. Notably, SPAdes correctly predicted these interspecific differences in SSC content as a barbell genome architecture for *P. antipodarum* and a lollipop genome architecture for *P. estuarinus*. We used a combination of Illumina MiSeq paired-end reads (IRs) and PacBio Circular Consensus Sequencing Reads (dinucleotide repeats) to insert the structural elements into each respective assembly, resulting in a closed, circular assembly in each case.

Both *de novo* assemblies exhibit consistent short-read mapping depth throughout the LSC (*P. antipodarum* mean coverage depth [+/- SD] = 879.83 [+/-95.17]; *P. estuarinus* mean coverage depth [+/-SD] = 5220.69 [+/-317.94]), and gene order is identical between the two species (Figure 2). Similarly, consistent coverage is obtained from mapping PacBio reads to the mitochondrial assemblies (Figure S2 a-b). When the nuclear genome assembly (BioProject ID: PRJNA717745) is excluded, coverage in the SSC spikes to >700,000x (Figure S2 c-d), likely as a result of simple-sequence repeats mapping here that are part of the nuclear genome. When the nuclear genome is included as part of the reference during read mapping, the short-read coverage of the SSC declines dramatically (Figure 2). Despite the decline in coverage, we are confident that this IR-SSC-IR structure is indeed part of the mitochondrial genome because (1) numerous PacBio subreads from each species (78 from *P. antipodarum* and 42 from *P. estuarinus*) span across the entire ∼1.7kb structure into unique sequence in the LSC region on each side and (2) there are no reads in any of our HiSeq, MiSeq, or PacBio datasets that span between the two ends of the LSC region that do not also contain the IR-SSC-IR element.

**Figure 2.**
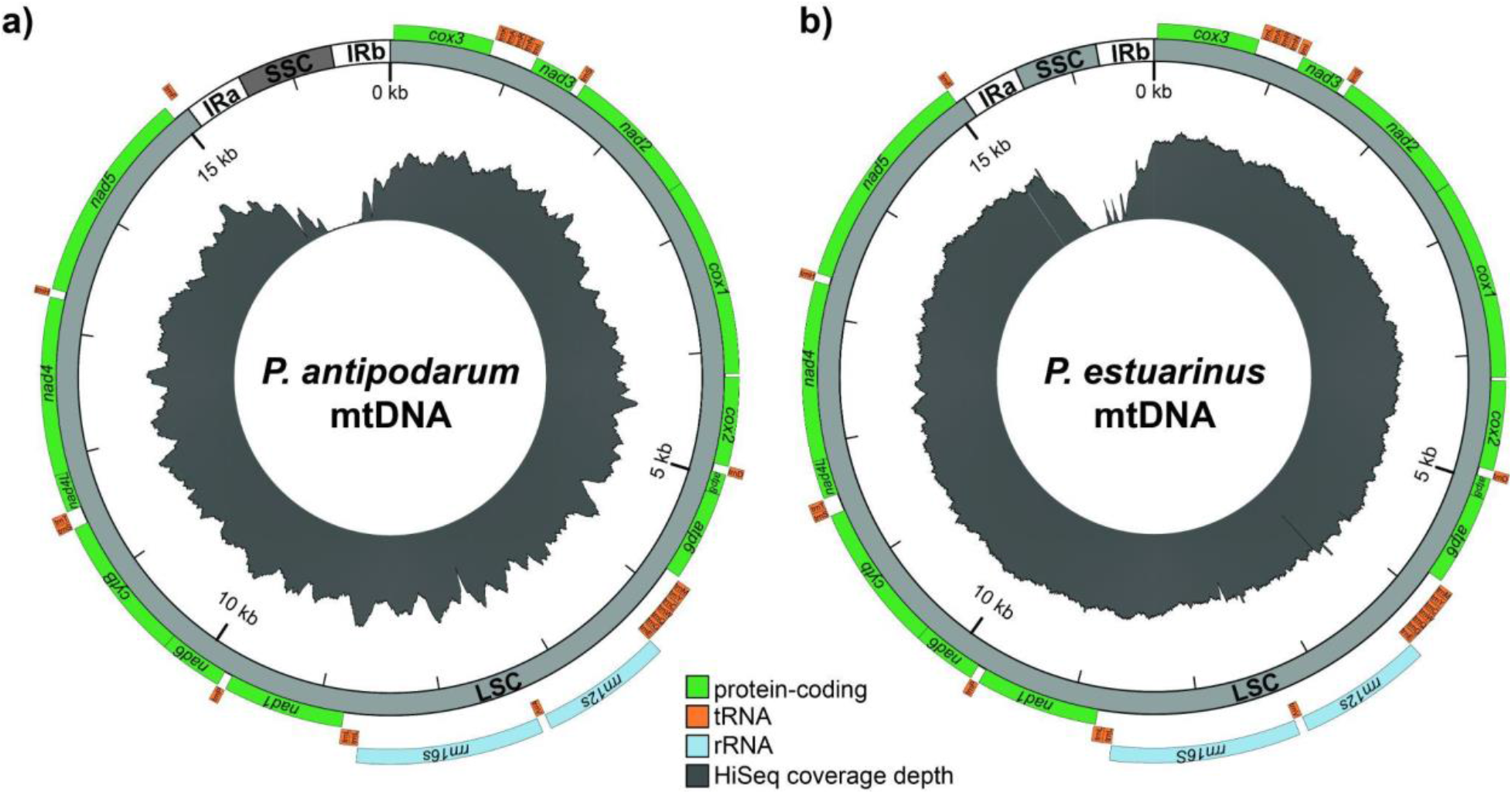
*Potamopyrgus* mitochondrial genome architecture. *De novo* mitochondrial genome assemblies depicted for a) *P. antipodarum* and b) *P. estuarinus*. Protein-coding (green), tRNA (orange), and rRNA (light blue) genes are arranged along a mitochondrial genome architecture consisting of a large single-copy region (LSC), two inverted repeats (IRa and IRb), and a small single-copy region comprised of dinucleotide repeats (SSC). Illumina HiSeq depth-of-coverage is depicted in the innermost circle in grey. The drops in coverage associated with the IRs likely reflect recombination breakpoints, while the low coverage in the SSC is likely due to low complexity reads mapping to non-mitochondrial contigs in the nuclear genome assembly.

We also confirmed the presence of the IRs by applying a custom-designed PCR approach using a single outward-facing primer paired with either a primer matching the top strand of the 5’ flanking sequence or the bottom strand of the 3’ flanking sequence (Table S1). That is, the primer designed to hybridize to the top strand of IRa (allowing polymerization towards *trnF* and *nad5*) can also anneal to the bottom strand of IRb (allowing polymerization towards *cox3*). The target amplicon spanning the *trnF*-IRa junction was designed to be 518 bp in length, while the amplicon targeting the IRb-*cox3* junction was designed to be 601 bp (Figure S3a). Both products were able to be amplified from both species (Figure S3b). We also attempted to amplify the SSC region using the reverse complement of the IR primer (*i.e*., anneals to the bottom strand of IRa and the top strand of IRb, potentially allowing for amplification of the SSC region). However, despite multiple attempts under the same conditions and multiple attempts with different conditions, including using a long-range polymerase and extension times up to six minutes, we were unable to recover an amplification product. It is possible that the high AT content in the SSC does not permit efficient amplification (consistent with previous unsuccessful efforts to sequence the SSC region with Sanger reads). Still, our PCR efforts directly confirm the presence of an inverted repeat independent of any sequencing data or library preparation artifacts.

Sliding-window analyses of patterns of substitution in *P. antipodarum vs*. *P. estuarinus* indicate that IRs exhibit high rates of sequence evolution relative to the rest of the genome (Fig. 3). The SSC also appears to evolve rapidly, as evidenced by the presence of a second dinucleotide repeat in *P. antipodarum vs*. only a single dinucleotide repeat in *P. estuarinus.* We also were able to determine that the inverted repeat is expressed from RNA sequencing reads. BLAST searches of the transcribed sequence did not have any significant hits, potentially indicating that the IR produces a novel non-coding RNA (*e.g.*, (53)). A more detailed survey of mitochondrial genome architectural diversity in molluscs and a broader sample of bilaterians may benefit tremendously from the wide availability of long-read sequencing data.

**Figure 3.**
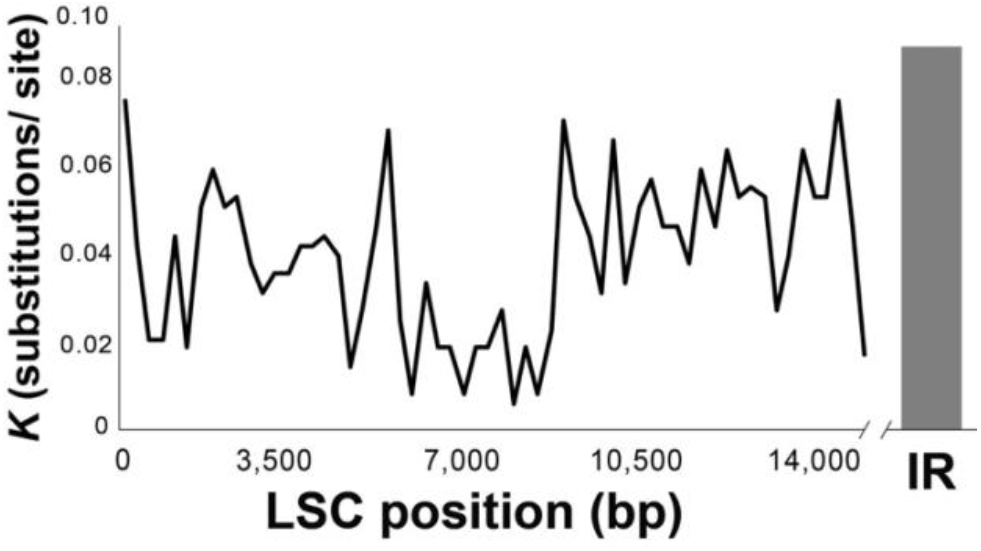
Sliding-window analysis of substitution rate (*K*) in the LSC (black line) compared to the single window of the inverted repeat (gray bar). Window size was 500 bp, with a 100bp step size.

Except for the IR-SSC-IR structure (Figure S4), we did not find any sequence differences between our assembly and the previous Sanger sequence-generated assembly for the mitochondrial genome of *P. antipodarum* (GenBank Acc. No.: MG979468.1). Evidently, 23 bp of the IRs nearest the LSC boundaries (46 bp in total) were used to circularize the old assembly. Other efforts to assemble this region in different *P. antipodarum* lineages (GQ996415-GQ996433, (25); MG979458-MG979470 (28)) took similar approaches.

The IR-SSC-IR architecture has profound implications for mitochondrial genome biology, particularly regarding mtDNA replication and repair. Indeed, IR-SSC-IR architectures are prone to forming hairpin or cruciform secondary structures, with the former implicated in DNA polymerase stalling (54). The predicted structure of IR-SSC-IR is somewhat reminiscent of the structure formed during vertebrate mtDNA replication at the light-chain replication origin (OriL), though the IR-SSC-IR is substantially longer (OriL ∼30 bp *vs*. IR-SSC-IR ∼1,700 bp). In vertebrates, mtDNA replication generally involves a strand-displacement mechanism in which leading-strand synthesis proceeds until encountering OriL, a ∼30-bp region that forms a stem-loop structure when single stranded. Lagging-strand synthesis is then primed starting in the loop of the single-stranded stem-loop structure (55, 56). Whether this genomic structural feature is involved in mtDNA replication represents an open question in *P. antipodarum*. However, the symmetrical orientation of the IRs raises the question of whether replication could proceed in both directions (either *via* displacement synthesis (15) or *via* strand-coupled synthesis (57)). Because mtDNA replication accuracy likely plays a central role in driving the notably high rate of mtDNA evolution in bilaterians, additional investigation into the diversity and distribution of mtDNA replication mechanisms will be critical for understanding mitochondrial genome biology and evolution.

### Repeat-associated recombination in Potamopyrgus mitochondrial genomes

Our observation that *Potamopyrgus* mitochondrial genome architecture resembles that found in chloroplast genomes of land plants (58) suggests that intra-molecular recombination between the IRs may lead to “flip-flopping” of the SSC, as originally suggested for chloroplast genomes by (51). While the SSC of *P. estuarinus* is itself an inverted repeat (*i.e.*, TA[x334]), and therefore impossible to use to determine directionality, the SSC of *P. antipodarum* (*i.e*., TA[x54]-TC[x337]) has definitive directionality. If flip-flopping is occurring, both orientations of the SSC should be detectable among PacBio reads spanning the region (Figure 4a). Consistent with this expectation, among 131 PacBio subreads that mapped unequivocally to both the LSC and the SSC in *P. antipodarum*, 77 subreads (58.8%) exhibit the forward orientation of the SSC, and 54 subreads (41.2%) exhibit the reverse orientation (Figure 4c). We also found that among 20 polymerase reads with multiple subreads mapping to both the LSC and the SSC, consecutive subreads invariably exhibited opposing orientations of the SSC after being oriented relative to the LSC (Figure S5). We interpret this invariable alternation of SSC orientation across consecutive subreads as evidence that the different strands of the mitochondrial genome have different SSC orientations, potentially indicating the presence of mitochondrial dimers. Several potential molecular mechanisms may explain this observation (see additional discussion below), all of which require some form of intra- or inter-molecular recombination. Together, these data provide strong and direct evidence of flip-flopping of the SSC relative to the LSC regions by frequent intramolecular recombination between the IRs. These data also explain why assembling an animal mitochondrial genome with only Illumina HiSeq data was so challenging.

**Figure 4.**
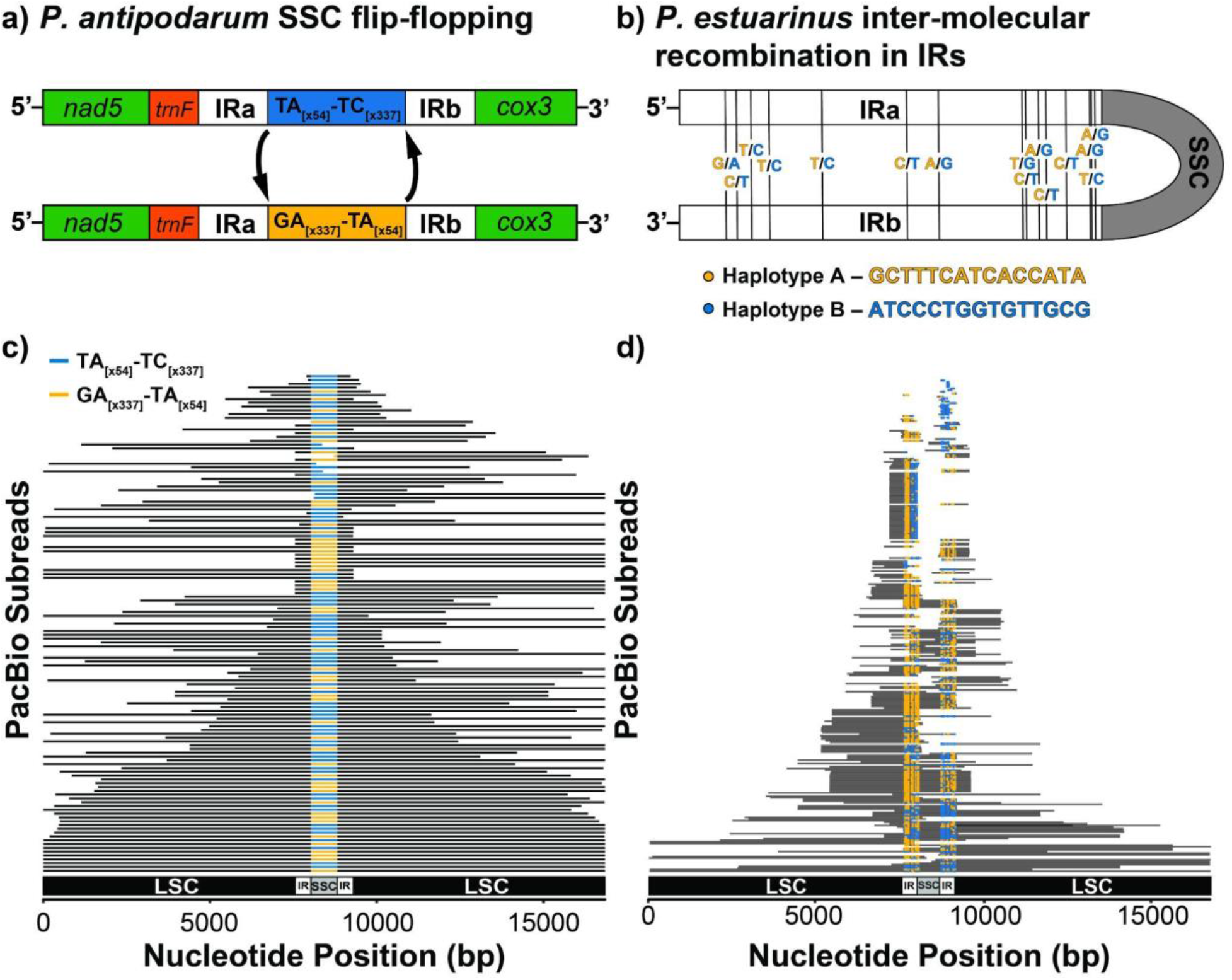
Repeat-mediated recombination in mitochondrial genomes of *Potamopyrgus*. a) Flip-flop recombination results in forward (top) and reverse (bottom) orientations of the SSC relative to the LSC in *P. antipodarum*. b) Two IR haplotypes exist in *P. estuarinus* mitochondrial reads, suggestive of inter-molecular recombination. c) Forward (blue) and reverse (orange) orientations of the SSC relative to the LSC exist at high frequencies among PacBio subreads from *P. antipodarum*. d) Sites of inter-molecular recombination in PacBio subreads from *P. estuarinus*.

Our long-read sequencing data provide useful glimpses into recombination mechanisms that could generate these patterns of flip-flopping. In particular, reads containing forward-oriented SSCs (*i.e*., matching the assembly presented herein) are not significantly more common than reads containing the reverse-oriented SSC (χ^2^ = 2.02, *p* = 0.155). This pattern is consistent with IR-mediated recombination equilibrium dynamics. Alternatively, the strict alternation of SSC orientations observed in *P. antipodarum* would seem to conflict with flip-flopped orientations resulting from equilibrium dynamics of IR-mediated recombination. This observation leads us to speculate that mitochondrial dimers, joined at the IRs, might explain the strand-specific nature of the flip-flopping. To assess whether we had captured mitochondrial dimers in our dataset, we identified 854 subreads from *P. antipodarum* that mapped to the mitochondrial assembly but that were longer than the length of the assembly. Of these 854 subreads, nine (1.1%) subreads (originating from seven distinct polymerase reads) from *P. antipodarum* appeared to capture an additional mtDNA molecule (Figure S6). There was one subread that bridged across the SSC multiple times, and for which the orientations were in the same direction in that case (*i.e*., m54138_171231_122620/22610103/17646_34950 TA-TC). Together, these data point towards IR-mediated recombination, resulting in the constant flip-flopping of the SSC. This same phenomenon is commonly observed in plant chloroplast genomes that harbor an inverted repeat (59).

Mitochondrial genomes, unlike chloroplast genomes that reside inside organelles that rarely fuse (60, 61), often encounter one another inside the cell during the process of mitochondrial fusion (62). Indeed, mitochondrial fusion and subsequent recombination among mtDNA molecules is thought to be a primary mechanism for maintaining mtDNA integrity in the face of oxidative damage (63). The widespread prevalence of mitochondrial fusion in eukaryotes combined with our observation of extensive flip-flop recombination in *P. antipodarum* mitochondrial genomes led us to speculate that *Potamopyrgus* mitochondrial genomes might also experience inter-molecular mitochondrial recombination associated with the IRs. We did not find any polymorphisms within the IRs with which we might map inter-molecular recombination, likely because the lineage we used for our reference genome sequencing has been inbred for more than 20 generations. Fortunately, *P. estuarinus* has no such inbreeding history, and all *P. estuarinus* individuals sequenced during this project were collected from the wild in New Zealand.

To test for inter-molecular recombination in *P. estuarinus* mitochondrial genomes, we first identified 15 high-frequency SNPs present in both Illumina MiSeq and PacBio read datasets generated from pools of multiple wild-caught *P. estuarinus* females (Figure 4b). We originally suspected these SNPs to represent differences among IRs rather than any form of heteroplasmy, particularly in light of our hypothesis that this region represents the hyper-variable D-loop. However, most PacBio reads that could span across both IRs harbor the same allele at both nucleotide positions (Table 1), indicating that the SNPs most likely come from different mitochondrial haplotypes rather than representing the accumulation of divergence between IRs. We used these same PacBio reads to phase the SNPs into two distinct haplotypes, A and B (Figure 4b). While haplotype A appears to be more common among this read pool than haplotype B (Table 1), there are many individual reads that harbor SNPs from both A and B haplotypes (Figure 4d). In sum, while there is little variation between the IR copies within a single mtDNA molecule, there are many combinations of alleles among distinct mtDNA molecules. This pattern implies that both intra- and inter-molecular recombination contribute to the distribution of genetic diversity in *Potamopyrgus* mtDNA.

**Table 1.** Inverted repeat haplotype frequencies in *P. estuarinus* PacBio pool-seq.

| IRa<br>Position | IRb<br>Position | Haplotype A<br>Allele | Haplotype B<br>Allele | A-A<sup>1</sup> (%) | A-B<sup>2</sup> (%) | B-A<sup>3</sup> (%) | B-B<sup>4</sup> (%) |
| --- | --- | --- | --- | --- | --- | --- | --- |
| 7603 | 9098 | G | A | 27 (47 %) | 7 (12 %) | 0 (0 %) | 23 (40 %) |
| 7615 | 9086 | C | T | 23 (44 %) | 13 (25 %) | 0 (0 %) | 16 (31 %) |
| 7631 | 9070 | T | C | 29 (64 %) | 5 (11 %) | 0 (0 %) | 11 (24 %) |
| 7651 | 9050 | T | C | 34 (61 %) | 8 (14 %) | 1 (2 %) | 13 (23 %) |
| 7709 | 8992 | T | C | 37 (66 %) | 7 (13 %) | 0 (0 %) | 12 (21 %) |
| 7802 | 8899 | C | T | 21 (35 %) | 18 (30 %) | 1 (2 %) | 20 (33 %) |
| 7837 | 8864 | A | G | 26 (43 %) | 12 (20 %) | 0 (0 %) | 22 (37 %) |
| 7929 | 8772 | T | G | 35 (48 %) | 13 (18 %) | 2 (3 %) | 23 (32 %) |
| 7933 | 8768 | C | T | 38 (54 %) | 10 (14 %) | 0 (0 %) | 23 (32 %) |
| 7947 | 8754 | A | G | 31 (47 %) | 8 (12 %) | 0 (0 %) | 27 (41 %) |
| 7955 | 8746 | C | T | 22 (32 %) | 7 (10 %) | 0 (0 %) | 39 (57 %) |
| 7977 | 8724 | C | T | 32 (44 %) | 20 (27 %) | 0 (0 %) | 21 (29 %) |
| 8003 | 8698 | A | G | 29 (48 %) | 16 (27 %) | 0 (0 %) | 15 (25 %) |
| 8005 | 8696 | T | C | 29 (48 %) | 17 (28 %) | 0 (0 %) | 15 (25 %) |
| 8009 | 8692 | A | G | 30 (48 %) | 18 (27 %) | 0 (0 %) | 18 (27 %) |

<sup>1</sup> – Number (percentage) of reads homozygous for the Haplotype A allele across the IR.
<sup>2</sup> – Number (percentage) of reads homozygous for the Haplotype A allele in IRa and the Haplotype B allele in IRb.
<sup>3</sup> – Number (percentage) of reads homozygous for the Haplotype B allele in IRa and the Haplotype A allele in IRb.
<sup>4</sup> – Number (percentage) of reads homozygous for the Haplotype B allele across the IR.

To gain a broader perspective of recombinational activity in *Potamopyrgus* mtDNA beyond the IRs, we also identified SNPs present in the LSC from the pooled *P. estuarinus* MiSeq data. In total, we identified 264 sites that were variable within the pooled sequencing (Table S2, Figure 5). After mapping MiSeq reads to the LSC, we combined paired-end reads into super reads using mapping information and quantified recombination between all pairs of SNP sites in the LSC. For each pair of SNP positions, we determined the haplotype of each read covering both sites and quantified the number of reads supporting each of the four combinations of alleles (*i.e*., REF-REF, REF-ALT, ALT-REF, and ALT-ALT). There were 4,697 SNP pairs with at least a single read covering both positions and 2,701 SNP pairs with >90x coverage (∼5-fold higher than expected nuclear coverage assuming mean insert size ≥ 600 bp). In this case, the presence of more than two haplotypic classes for any given SNP pair was used as evidence of the activity of inter-molecular recombination. Among the 4,697 SNP pairs, we found 367 pairs with >90 reads supporting the third-most-common haplotypic class. We even found 32 SNP pairs with >180 reads covering the third-most-common haplotypic class. There were fewer pairs in which the fourth-most-common haplotypic class exceeded the 5x threshold expected of the nuclear genome, as only two pairs were found in which the fourth most common haplotypic class met or exceeded the 90x coverage threshold. When only considering the 263 independent pairs of SNPs (*i.e.*, adjacent SNPs), we found that 139 pairs (52.9%) exhibited >90x coverage of the third-most-common haplotypic class (Figure 5, Table S3). Conversely, only 125 (46.4%) of the adjacent SNP pairs exhibited <90x sequencing coverage over the 3rd most common haplotype class, indicating that recombination between adjacent SNPs was more likely than not. One of the adjacent SNP pairs (10573 [C/T] – 10630 [T/C]) exhibited relatively high coverage of all four haplotype classes, with 365 C-T reads, 78 T-T reads, 243 C-C reads, and 312 T-C reads. In sum, the high frequency of SNP pairs with >2 haplotypic classes at coverage levels far exceeding those expected by nuclear-encoded mitochondrial sequences (numts) or that can be explained by PCR or sequencing errors supports the presence of inter-molecular recombination in *P. estuarinus* individuals.

**Figure 5.**
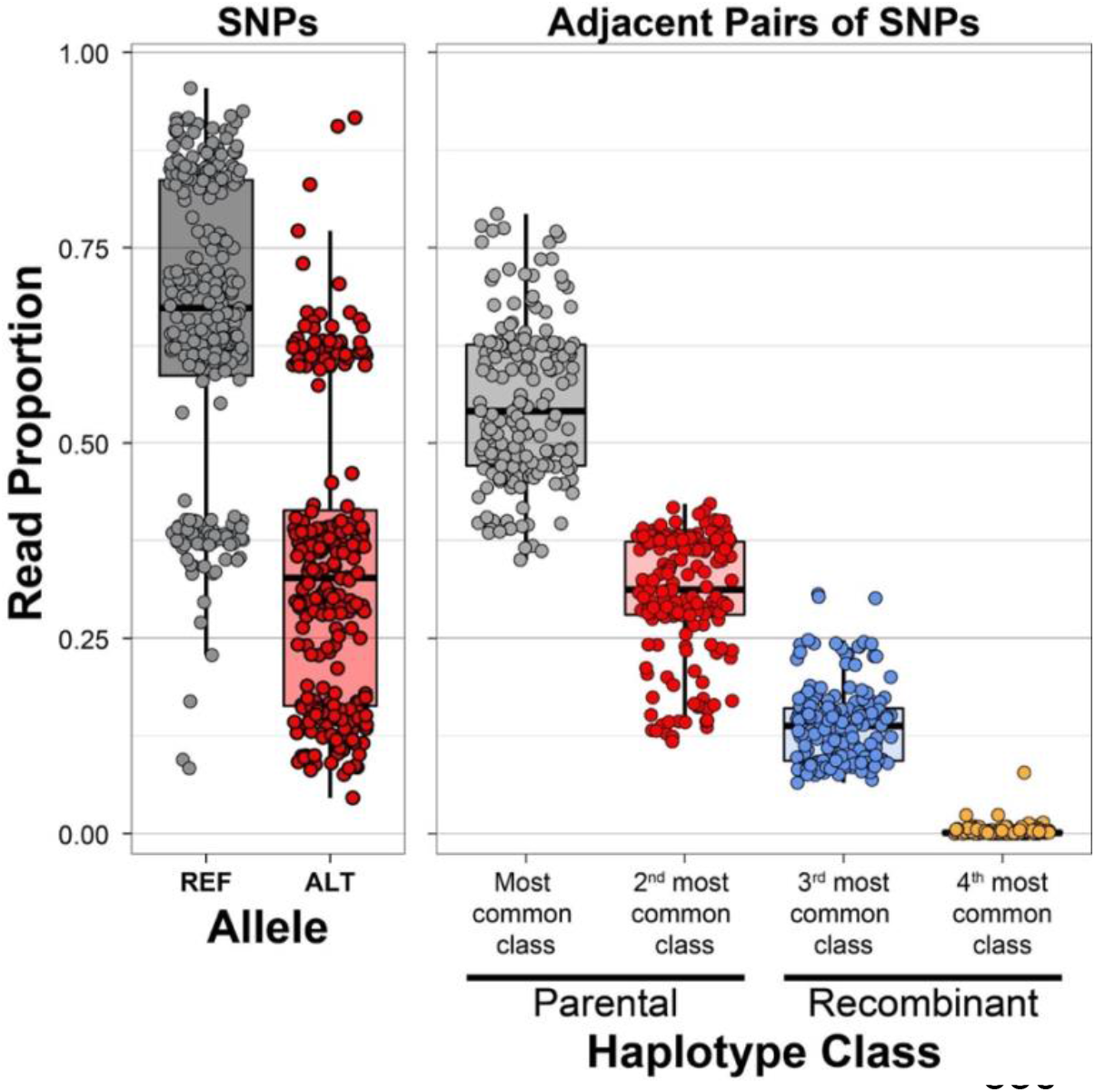
Polymorphism and recombination in *P. estuarinus* mitochondrial genomes. Left – Allele frequencies for individual SNPs. Minimum read depth at SNP positions was 349x, with at least 47x coverage over minor alleles at all positions. Right – Haplotype class frequencies for pairs of SNPs measured by Illumina MiSeq read depth. The presence of more than two haplotype classes for any given pair of SNPs implies recombination among the molecules sequenced. Of the 263 independent SNP pairs considered here, 182 exhibited >69x sequencing coverage over the third most common haplotype class.

The presence of inter-molecular recombination documented above implies extensive heteroplasmy within individual cells, as mtDNA molecules must co-exist inside the same cell (indeed, the same mitochondrion) in order to recombine. While these data do not provide direct evidence regarding the mode of mitochondrial inheritance, we speculate that the apparent large degree of heteroplasmy required to explain these data is likely to have originated from multiple parents (13). Perhaps even more surprising is the extensive mitochondrial diversity maintained in *P. estuarinus* populations in the face of recombination. These snails were collected within meters from one another, yet hundreds of mitochondrial SNPs can be recovered from only eight individuals. The mechanism(s) whereby such high levels of mitochondrial diversity are maintained in this system represent an important area of future investigation.

Mitochondrial recombination has been detected in a wide diversity of plant, fungi, and even animal taxa (12, 64–72). Indeed, plant geneticists have long recognized the importance of recombination for mitochondrial genome architecture and evolution (73, 74). A recent study even reported a history of extensive genetic recombination in the massive mitochondrial genomes of populations of the angiosperm *Silene* (75). More commonly, plant mitochondrial genomes undergo extensive repeat-mediated recombination. So-called “substoichiometric shifting” (76–79) results in multiple distinct mitochondrial isoforms existing within the same cell (80, 81). Fungal mitochondrial genomes also appear to experience relatively frequent homologous recombination (71, 82–85).

By contrast, evidence for mitochondrial recombination in animals is relatively scarce (86) and has primarily been observed in comparative and population genetic datasets (87–91), with a couple of notable exceptions. One prominent example is provided by Doubly Uniparentally Inherited (DUI) mitochondrial genomes in some mollusc taxa, which can facilitate recombination (66, 92). Mitochondrial genomes in human and mouse heart tissue have been found to form recombination-associated structures during mitochondrial genome replication (93, 94), and intra- and inter-molecular recombination between distinct mitochondrial molecules in heteroplasmic individuals can be induced through oxidative stress in *Drosophila* (70) and using restriction enzymes in mice (95). Recent reports of heteroplasmy in humans (96, 97) underscore the potential importance of mitochondrial recombination to human health outcomes.

Repeat-mediated recombination in plant mitochondrial genomes is strongly associated with DNA repair *via MSH1* and is often cited as a primary cause of their relatively low rate of molecular evolution (79, 98, 99). By contrast, the high levels of nucleotide diversity and high rates of sequence evolution in the *Potamopyrgus* IRs suggest that recombination is not acting to reduce the rate of sequence evolution. The recombination we observed in the LSC and the IR has two primary consequences. First, recombination between distinct mtDNA molecules can reset Muller’s ratchet (100, 101), which is otherwise expected to lead to the eventual collapse of asexual lineages (102). Second, recombination between distinct mtDNA molecules breaks down linkage disequilibrium between mutations in the mitochondrial genome, countering the Hill-Robertson effect (103) by allowing these mutations to be selected (positively or negatively) independently from their mitochondrial genomic background. Under the presumption that mitochondrial genomes lack recombination, both Muller’s ratchet and the Hill-Robertson effect have long been assumed to generate harmful mutation accumulation in mitochondrial genomes (101). However, recent work in nematodes (104) and other animals (105) indicates that the effective population size of mtDNA may be much larger than previously thought (106, 107). Our discovery of inter-molecular recombination in these molluscs and the continued existence of separately inherited nuclear and mitochondrial genomes after more than a billion years of intra-cellular co-evolution are consistent with the hypothesis that mitochondrial recombination might facilitate escape from seemingly inevitable mutational meltdown.

### Summary

We document a novel and complex genomic structural feature of the mitochondrial genomes of *Potamopyrgus* snails. These native New Zealand prosobranch snails are well studied because of the status of *P. antipodarum* as a model system for sexual reproduction, host-parasite coevolution, and as a worldwide invader. Interest in how the consequences of reproductive mode polymorphism might influence mitochondrial evolution and the maintenance of sex has motivated surveys revealing extensive genetic variation in mtDNA (25, 28, 108, 109) as well as phenotypic variation for mitochondrial function (39, 40) in *P. antipodarum*. While previous efforts to sequence *P. antipodarum* mitochondrial genomes have provided valuable data on this front (25, 28), all have been restricted to short sequencing reads or PCR amplified segments, and none have been able to assemble the entire circular structure.

Using a combination of short (Illumina HiSeq, Illumina MiSeq) and long (PacBio) reads obtained from the initial stages of sequencing the whole nuclear genome, we successfully re-assembled the entire *P. antipodarum* and *P. estuarinus* mitochondrial genomes *de novo*. In addition to confirming the presence of the previously reported sequence (25, 28), these assemblies revealed a pair of inverted repeats (IRs) interrupted by a repetitive single-copy region that were not recognized in the earlier studies. The IRs appear to evolve very rapidly (in contrast to the slow rates of molecular evolution seen in chloroplast IRs (110)), and RNAseq data revealed the presence of a novel transcript, ∼450 bp in length, with no open reading frames or identified homology. The nature of this genomic feature and its clear resemblance to the genomic architecture of plant chloroplast genomes (11) suggests that this component of *Potamopyrgus* mitochondrial genomes may be a site for intra- and inter-molecular repeat-mediated recombination. PacBio reads confirmed this hypothesis and indicated that intra-molecular recombination resulted in “flip-flopping” (51) of the small single-copy region (SSC) relative to the large single-copy region (LSC) is very frequent and that inter-molecular recombination between distinct mtDNA molecules is common when present within the same cell. Together, these findings necessitate a careful re-examination of bilaterian mitochondrial genomes, with special emphasis on repeat-mediated mitochondrial recombination. More broadly, the potential for the widespread presence of recombination in bilaterian animal mitochondrial genomes that this result suggests will enable important new insights into mitochondrial biology and evolution and raise important questions: Are hyper-variable regions like D-loops recombination hotspots in mitochondrial genomes? Does mitochondrial recombination stave off mutational meltdown? Is the IR-SSC-IR genome architecture a cause or consequence of mitochondrial recombination?

## Materials and Methods

### DNA sequence data

DNA from *P. antipodarum* and *P. estuarinus* were sequenced as part of the *Potamopyrgus* nuclear genomes project using Illumina HiSeq, Illumina MiSeq, and PacBio SMRT sequencing (PRJNA717745). DNA extraction details and library preparation details are fully described elsewhere. Briefly, we extracted DNA from Alex Yellow (a sexual *P. antipodarum* lineage inbred for 20+ years) and from field-collected *P. estuarinus* using a guanidinium-thiocyanate-based phenol-chloroform DNA extraction protocol (28, 111), cleaned DNA extractions with Zymo’s Clean and Concentrate Kit (Zymo Research, Irvine, CA, USA), and re-suspended DNA in 30-100 µL of T-low-E buffer (10 mM Tris pH 8.0, 0.1 mM EDTA). DNA quality was assessed on a NanoDrop 1000 to determine DNA concentration and purity, and only DNA extractions that contained 200-1000 ng DNA and for which the 260/280 was ∼1.8 were used for library construction. 100 ng of each sample was run on a 0.8% agarose gel to confirm the integrity of the sample and to identify any obvious contaminants. We sequenced 2x100 nt paired-end reads on an Illumina Hi-Seq 2500 and 2x250 nt paired-end reads on an Illumina MiSeq. To obtain high-molecular weight DNA for PacBio sequencing, we extracted DNA from pools of Alex Yellow (N = 20) and *P. estuarinus* (N = 20) individuals with a user-developed protocol for Qiagen genomic tip DNA extraction for insects (Genomic-tip 100/G) that uses a gravity column for high-molecular weight extractions (Alex Yellow) and the protocol described in (112) (*P. estuarinus,* conducted at the Arizona Genomics Institute). The Pacbio sequencing library construction followed the PacBio 20-kb protocol in both cases. These libraries were sequenced in nine SMRT cells of a Sequel for Alex Yellow and 10 SMRT cells for *P. estuarinus*. All PacBio sequencing was performed at the Arizona Genomics Institute (Tuscon, AZ, USA), which provided all subread BAM files and associated SMRT movie metadata.

We evaluated Illumina DNA sequence data quality using the FASTQC program in the FASTX-Toolkit (113), trimmed adapters with cutadapt v.3.4, and trimmed low-quality bases (*i.e*., PHRED scores <20) with the FASTQ Quality Trimmer. PacBio Circular Consensus Sequencing (CCS) reads were obtained using ccs v3.4.0 with default settings (https://ccs.how/), requiring a minimum of 3 subread passes for CCS generation.

### Mitochondrial genome assembly

We assembled the paired-end HiSeq reads *de novo* using SPAdes v. 3.13.0 (31) separately for Alex Yellow and *P. estuarinus* with default settings. We then extracted mitochondrial contigs from the assembled contigs of each species using blastn v. 2.7.1+ (114) against the published mitochondrial genome from Alex Yellow (MG979468.1), setting the e-value threshold to 10^-5^ and max_target_seqs to 10,000. Significant blast hits were manually stitched together according to the published Alex Yellow genome assembly (25). We used Pilon v. 1.23 (52) to polish the mitochondrial assemblies with lineage-specific paired-end reads.

At this stage, both the Alex Yellow and *P. estuarinus* mitochondrial assemblies displayed a single discontinuity (*i.e*., position preventing circularization), leading us to hypothesize that a repeat element was causing the assembly to break at position 12,481 relative to the published Alex Yellow mitochondrial reference genome. We did not detect any other differences between our newly generated Alex Yellow mitochondrial assembly and the previously published assembly. Further investigation of the discontinuity revealed that the published Alex Yellow mitochondrial reference genome contained a 44-bp palindromic repeat at this site, potentially explaining the anomaly. However, paired-end Illumina reads could not span the structure, despite being of adequate length (*i.e*., >44 bp) to do so. To further investigate this discrepancy, we used Bandage v. 0.8.1 (115) to visualize the assembly graph (Figure S1), which revealed a clear signature of inverted repeats. As a result, we inferred that single-molecule PacBio data generated during the *P. antipodarum* and *P. estuarinus* nuclear genome sequencing projects would best be able to resolve the apparent discontinuity in these short-read assemblies.

First, we re-oriented our short-read assemblies to place the discontinuity at the center of the Alex Yellow and *P. estuarinus* assemblies. We then mapped PacBio subreads from each species to these mitochondrial assemblies using blasr v. 5.3.2-06c9543 (116). We collected reads that spanned the discontinuity, mapped the reads to themselves using minimap2 v 2.15-r905 (117), and assembled only those PacBio reads using miniasm v 0.3-r179 (118). We aligned the resulting long-read assembly to the mitochondrial assemblies of both species, which both contained a ∼1.7 kb previously undetected region that coincided exactly with the discontinuity. A dot plot of the structures revealed that their 5’ and 3’ flanks were inverted repeats (IRs), with an interior region comprised of dinucleotide repeats. In Alex Yellow, there were two distinct dinucleotide repeats (AT and CT), while only the AT repeat was present in *P. estuarinus*. We therefore inserted this IR-dinucleotide-IR structure into each of our respective short-read assemblies and used PacBio reads that mapped outside the discontinuity to polish the assemblies with Arrow v2.2.2 (119).

To confidently assemble the IRs, we mapped MiSeq reads to the mitochondrial genome assemblies and collected all reads that mapped within 100 bp of the 5’ end of the structure and within 100 bp of the 3’ end of the structure. We discarded reads whose alignments did not extend *beyond* the boundaries of the structure to ensure that we only included reads of clear mitochondrial origin. We assembled these MiSeq reads for the 5’ IR and the 3’ IR separately using SPAdes. In Alex Yellow, both 5’ and 3’ assemblies yielded a single identical contig that captured the entire length of the IR (*i.e*., spanned from the 5’ end to the dinucleotide repeat and from the 3’ end to the dinucleotide repeat). However, in *P. estuarinus*, 5’ and 3’ IR assemblies contained two distinct contigs. From these results, we inferred that the IRs contained 15 single nucleotide differences (see Figure 4b), which were later verified with PacBio CCS reads from *P. estuarinus* obtained from distinct sample collections and DNA extractions.

Finally, to assemble the dinucleotide repeats, we aligned PacBio CCS reads to the HiSeq-PacBio-MiSeq assemblies for each species using blasr and obtained consensus lengths for the two dinucleotide repeats in Alex Yellow (TA[x54]-TC[x337]) and the single dinucleotide repeat in *P. estuarinus* (TA[x334]). We validated our assemblies using 1) long single-molecule PacBio reads that could span the insertion and the artificially introduced assembly break at the 5’ and 3’ ends of the linearized assemblies, and 2) comparing coverage across the entire assembly using short reads.

We manually annotated genes from the previously published *Potamopyrgus* mitochondrial genome assemblies onto the newly generated assemblies, searched for novel ORFs (using the invertebrate mitochondrial genetic code) in the insertion using a blastp search against the nr/nt database as well against the *P. antipodarum* EST database (http://bioweb.biology.uiowa.edu/neiman/blastsearch.php) (120), and ran the assemblies through MITOS2 (121) to obtain tRNA structural predictions.

### PCR confirmation of inverted repeats

We designed a custom PCR approach to directly test the inverted repeat hypothesis using four experiments (Table S1). All PCR experiments were performed in 25 µL volumes, using 5 ng DNA, 2.5 µL 10X iTaq Buffer (Bio-Rad, Hercules, USA), 1.5 mM MgCl2, 200 µM for each dNTP, 0.1 µM of each primer, and 1U of iTaq DNA polymerase (Bio-Rad, Hercules, USA). The first experiment was a positive control to amplify a 240 bp region in the gene-dense LSC. The second experiment was designed to amplify a 518-bp fragment of the left flank of the inverted repeat using a forward primer complementary to the bottom strand of *trnF* and a reverse primer that was complementary to the top strand of IRa. For the third experiment, we used this same IR primer as a forward primer in which it had to be complementary to the bottom strand of IRb in order to amplify a 601-bp product in concert with a reverse primer designed to complement the top strand of *cox2*. Thus, the IR primer can be used to amplify two completely different regions of the mitochondrial genome, at the predicted sizes (Figure S3b). Finally, in an attempt to amplify the SSC region, we designed a final primer, which was the reverse complement of the IR primer. In this experiment, we used only a single primer, with experimental concentration doubled to 0.2 µM. The primer was predicted to be complementary to both the bottom strand of IRa and the top strand of IRb, and if amplification were successful, would produce an ∼1100-bp product. We used the same conditions for each PCR experiment: 95°C for a three-minute denaturation, then cycled the following three steps 40x: 1) 30 seconds at 95°C, 2) 30 seconds at 56°C for primer annealing, and 3) 1.5 minutes at 72°C for extension. We had a final extension time of six minutes and then evaluated the success of the PCR reactions on a 1% sodium borate-agarose gel. Because the fourth experiment did not yield any positive results, we also tried a six-minute extension time for each cycle, using a LongAmp polymerase (New England Biolabs, Ipswich, USA), which also did not yield any bands. Gels containing 3.5 µl of SYBR Safe stain (Invitrogen, Waltham, USA) were run at 300V for 15 minutes and visualized under blue or UV light.

### Identification of recombinant mitochondrial molecules

The resemblance of certain elements of the *Potamopyrgus* mitochondrial genome structure to that of chloroplast genomes in most land plants (11) led us to hypothesize that the IRs could mediate intra-molecular recombination. Because the SSC in Alex Yellow comprises two distinct dinucleotide repeats (*i.e*., TA and TC), reads that span the length of the IR-SSC-IR structure can readily be evaluated for recombination. If recombination exists in these mitochondrial genomes, both TA-TC *and* GA-TA SSC conformations should be observable among reads that span the structure. In *P. estuarinus*, the orientation of the SSC is not useful for diagnosing recombination because the SSC is identical in either direction. We therefore had to rely on 15 single-nucleotide differences between IRa and IRb (inferred from MiSeq reads and confirmed in PacBio CCS reads) to quantify rates and patterns of mitochondrial recombination in *P. estuarinus*.

We mapped PacBio subreads and CCS reads from each species to our final assemblies and quantified IR/SSC orientations in reads that 1) spanned the entire structure and 2) in all reads that overlapped with the IR-SSC-IR structure, but that also mapped unambiguously to one side of the LSC or the other. We visualized the orientations of these reads using CIRCOS v. 0.69-5 (122) and the overall read frequencies in R v4.1.2. We tested whether the forward orientation of the SSC was more common than the reverse orientation using a χ^2^ goodness-of-fit test.

Recombination among IRs in chloroplast genomes is known to reduce the rate of sequence evolution relative to rates in the single-copy regions (123). We therefore aligned IRs and LSCs separately and compared sequence divergence between *P. antipodarum* and *P. estuarinus* across the two regions, removing any sites containing polymorphism within *P. antipodarum* from the analysis. We excluded the SSC from this analysis because its repetitive nature makes for uncertain and dubious base calls and alignments.

Recombination between SNP pairs in the LSC was performed using MiSeq reads. We mapped MiSeq reads to the *P. estuarinus* assembly and used mapping information to convert paired-end reads into super reads using a custom python script (superReads.py). We re-aligned super reads to the mitochondrial assembly and converted the resulting SAM file to a multiple sequence alignment using a custom python script (sam2Fasta.py). Haplotype calls from individual reads were obtained from the multiple sequence alignment and quantified using the recombinantReads.py python script. All python scripts used in this study are available at (https://github.com/jsharbrough/potamoMitoGenomes).

### Mitochondrial transcriptomics

Taking advantage of existing RNA-seq dataset for both species, we determined whether *P. antipodarum* and *P. estuarinus* exhibit polycistronic transcription of their mitochondrial genomes and whether their mitochondrial transcripts experience RNA editing. Briefly, for both species, heads were removed from individual snails and used for flow cytometry to confirm diploidy. RNA was extracted using TRIzol from body tissue of snails from each of three conditions: one male, one non-brooding female, and one brooding female from the Alex Yellow *P. antipodarum* line, and one male, one reproductively active female, and one non-reproductively active female from field-collected *P. estuarinus* (*124*). Sequencing libraries were prepared using the Illumina Tru-Seq LS protocol, which were used for 2x100 bp paired-end Illumina HiSeq 2000 RNA sequencing (46). We used FASTQC to assess read quality (125), and FASTX Toolkit (113) to trim sequencing adapters and remove low-quality reads.

We employed two complementary methods to assess the presence of polycistronic transcripts. First, for each species and separately for each condition, filtered reads were mapped to their respective mitochondrial genome assemblies using Hisat2 (126) with --no-mixed and --no-discordant parameters set. Mapping was performed against two reference orientations to ensure that transcript assembly was not biased based on where the assembly was linearized: (1) LSC-IR-SSC-IR, and (2) LSC-IR-SSC-IR-LSC. We then used Samtools (127) to convert the hisat2 sam output into bam format, and then used PicardTools (128) to add read groups, remove PCR duplicates, and sort bam files according to coordinates. Next, transcripts were assembled using StringTie (129). Using the published mitochondrial genome annotations, we assessed the assembled transcripts for the presence of polycistronic transcripts.

Second, for both species and each condition, we used BedTools bamtofastq (130) to extract mapped reads from the processed bam files. Within condition, we concatenated the left and right read files (*e.g.,* from *P. antipodarum* males from the two reference orientations) and removed duplicated reads. We then used Trinity to generate *de novo* transcript assemblies for each species and for each of the three conditions. These transcripts were then compared to the established mitochondrial genome annotations and the transcripts generated by StringTie to further confirm the presence of polycistronic transcripts.

To evaluate whether *Potamopyrgus* mitochondrial transcripts experience RNA editing, we mapped the HiSeq genomic reads to the LSC-IR-SSC-IR orientation mitochondrial reference using Hisat2 and processed the bam files as stated above. We then used REDItools.py (131) to identify SNPs present in the RNA-Seq data for each species and condition that was not present in the genomic sequencing data (representing genetic variation as opposed to edited sites). We invoked REDItools parameters -n 0.05 -N 0.05 -e -E -d -D -u -U -L -v 2, and otherwise implemented default parameters.

### Data availability

All sequence data (genomic and transcriptomic) supporting the mitochondrial assemblies are available in Genbank under BioProject PRJNA717745. Mitochondrial assemblies are available in Genbank (Accession Nos. ####). Mitochondrial genome assemblies and annotations, sequencing reads, and transcriptome assemblies are also available on FigShare (*P. antipodarum* – https://doi.org/10.6084/m9.figshare.20471784; *P. estuarinus* – https://doi.org/10.6084/m9.figshare.20474262). All scripts are available at the following GitHub repository (https://github.com/jsharbrough/potamoMitoGenomes).

## Acknowledgements

We thank Cindy Toll for assistance with DNA extraction, and Gus Waneka, Dan Sloan, and Sam Ward for comments on the manuscript, discussions about mechanisms of recombination, and statistical design. We thank Cindy Toll, Katelyn Larkin, and Peter Wilton for contributions to the *Potamopyrgus* genome project. We acknowledge funding from the National Science Foundation (NSF-MCB 1122176, NSF-DEB 1753851), NM-INBRE, and New Mexico Institute of Mining and Technology. Some of this work utilized resources from the University of Colorado Boulder Research Computing Group, which is supported by the National Science Foundation (awards ACI-1532235 and ACI-1532236). Illumina sequencing was performed at Iowa Institute of Human Genetics (IIHG) Genomics Division, and PacBio sequencing was carried out at the Arizona Genomics Institute.

## SUPPLEMENTARY FIGURES

**Figure S1.**
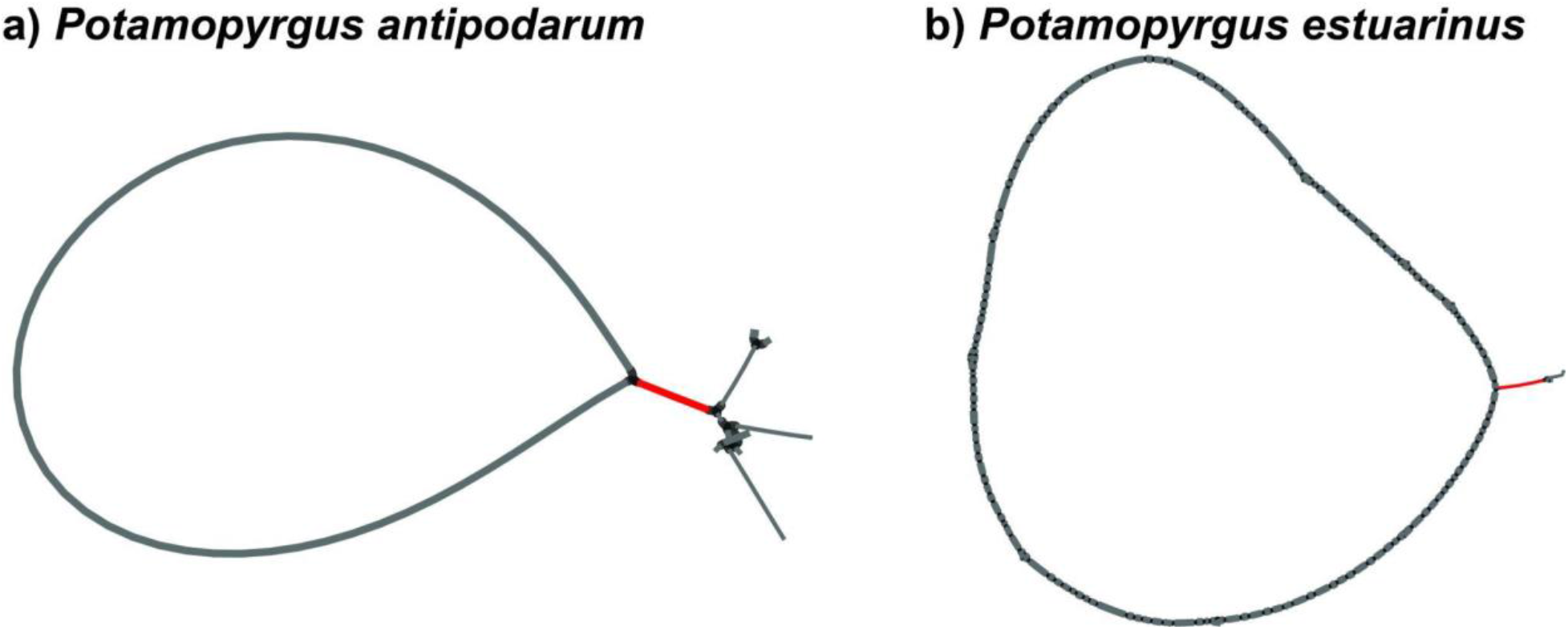
Visualizing *Potamopyrgus* genome architecture. a) Barbell architecture representing *P. antipodarum* mitochondrial contigs after short-read assembly. b) Lollipop architecture representing *P. estuarinus* mitochondrial contigs after short-read assembly. Barbells and lollipops in assembly graphs are characteristic of inverted repeats within circular genomes. Dot plots of the novel 1.7kb structures in c) *P. antipodarum* and d) *P. estuarinus* depict inverted repeats interspersed by a small single-copy region.

**Figure S2.**
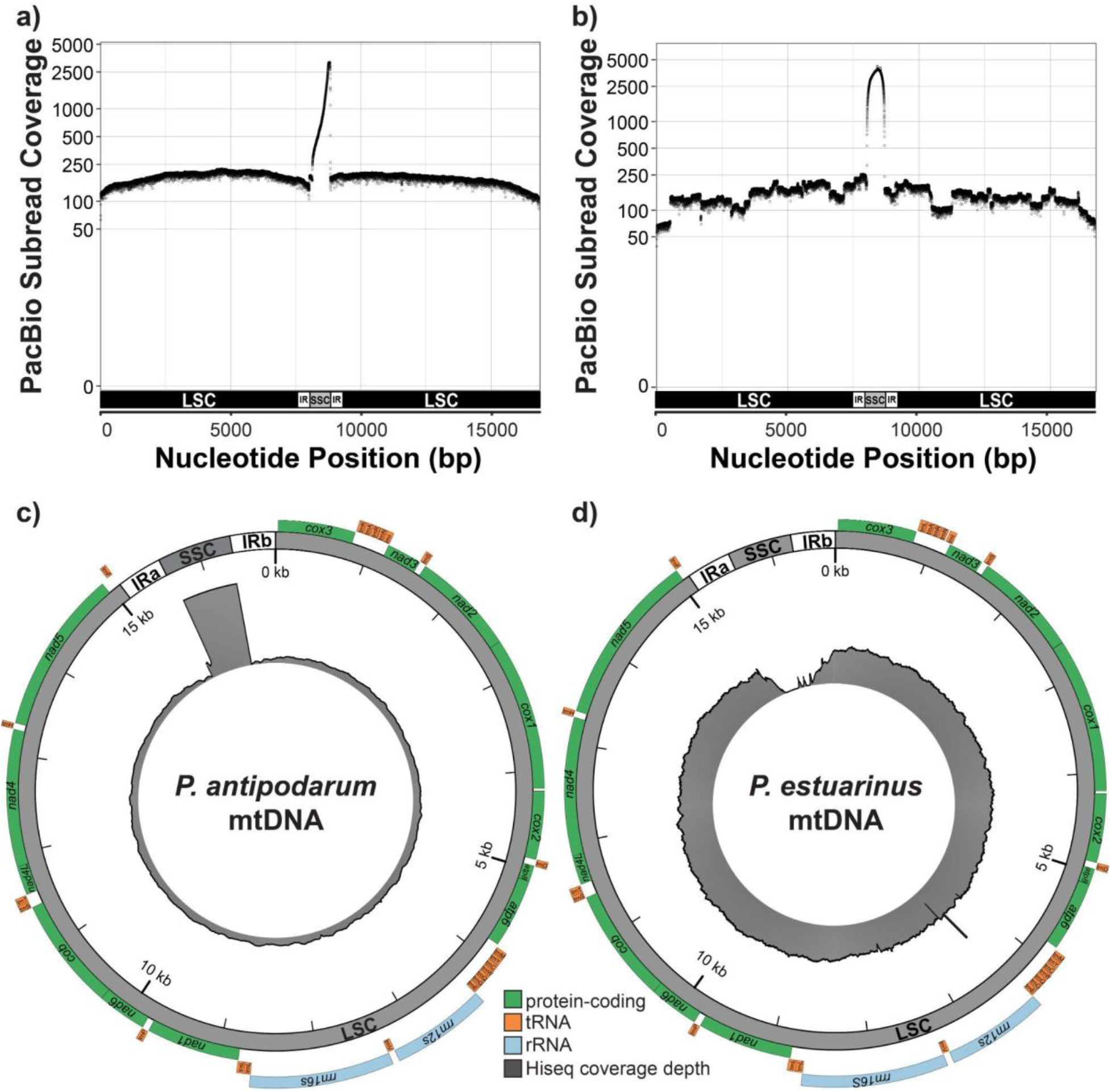
Pacbio and Illumina HiSeq coverage when the nuclear genome is excluded from the mapping experiment. a), c) Depth of coverage for *P. antipodarum*, and b),d) Depth of coverage for *P. estuarinus* mitochondrial genomes. a), b) PacBio read depth (black circles) calculated with mosdepth using default parameters. Log-base-10 read counts across the mitochondrial genome are shown for both species. c),d) Illumina HiSeq coverage depth (grey bars surrounding inner circle) when mapped to both species mitochondrial genomes.

**Figure S3.**
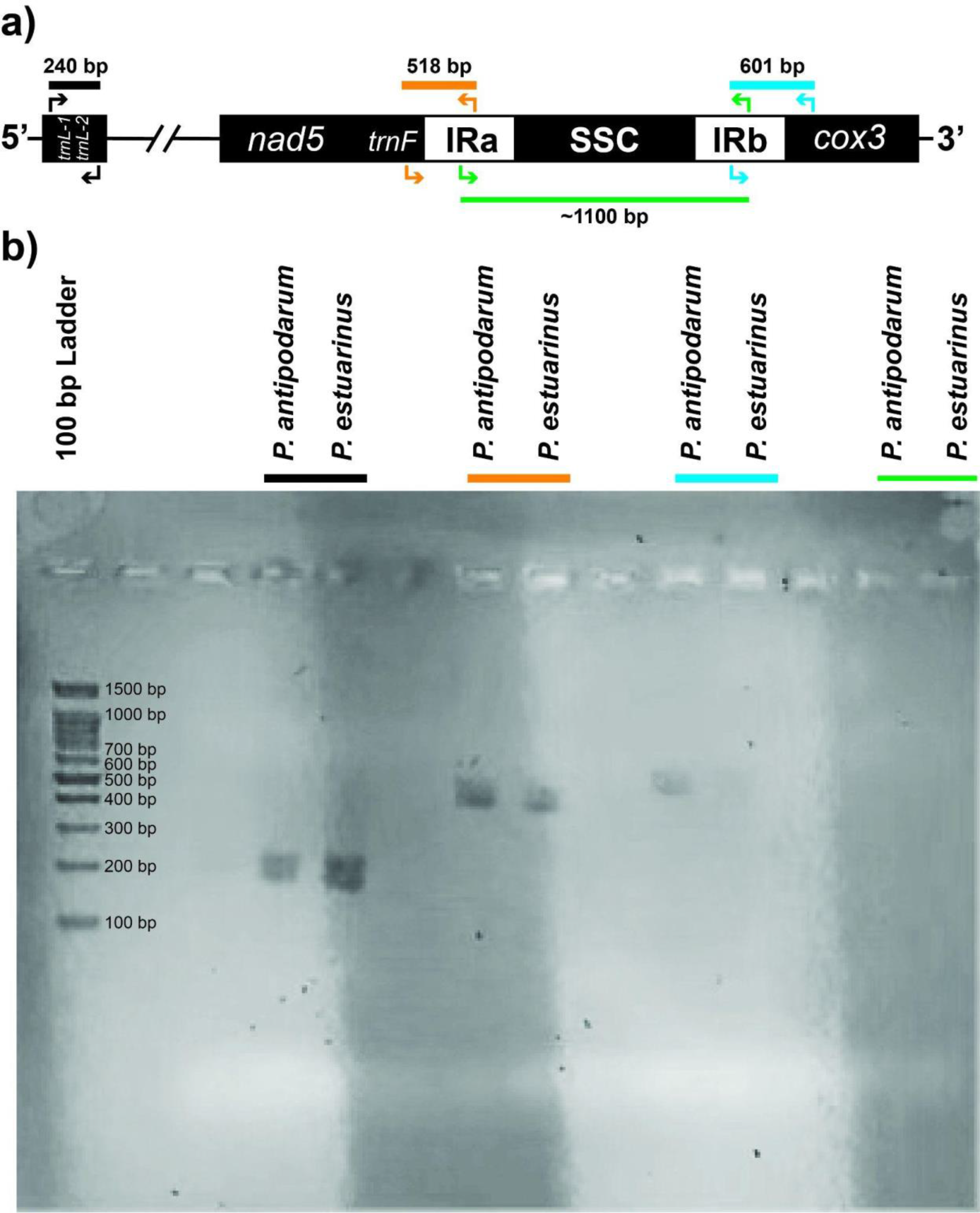
PCR confirmation of inverted repeats. a) Cartoon schematic depicting PCR experiment design. b) Example gel image taken featuring DNAs from both species, following PCR amplification for all four sets of primers. All primer pairs worked, with the exception of the SSC experiment (green bar).

**Figure S4**. Alignment of the newly assembled mtDNA from the inbred Yellow lineage from Lake Alexandrina, NZ compared to the published mitochondrial assembly (MG979468) from the same inbred *P. antipodarum* lineage. No differences in nucleotide sequence exist, except that the old assembly lacks the IR-SSC-IR structure (orange arrows). Instead, a 46 bp inverted repeat (underlined in red) present in MG979468 was used to circularize the assembly.

**Figure S5.**
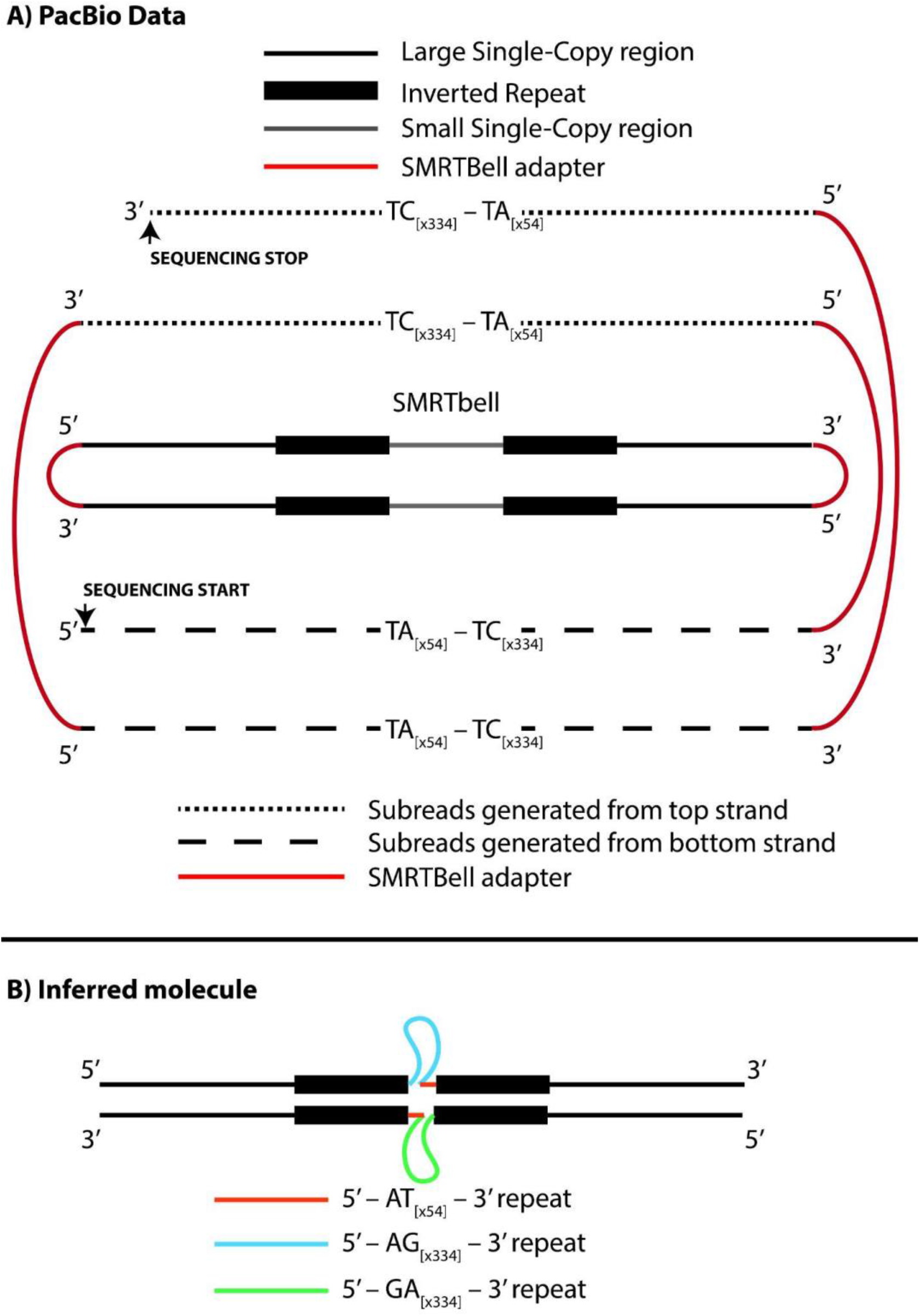
Cartoon depicting observed alternation of SSC orientation from consecutive subreads.

**Figure S6.**
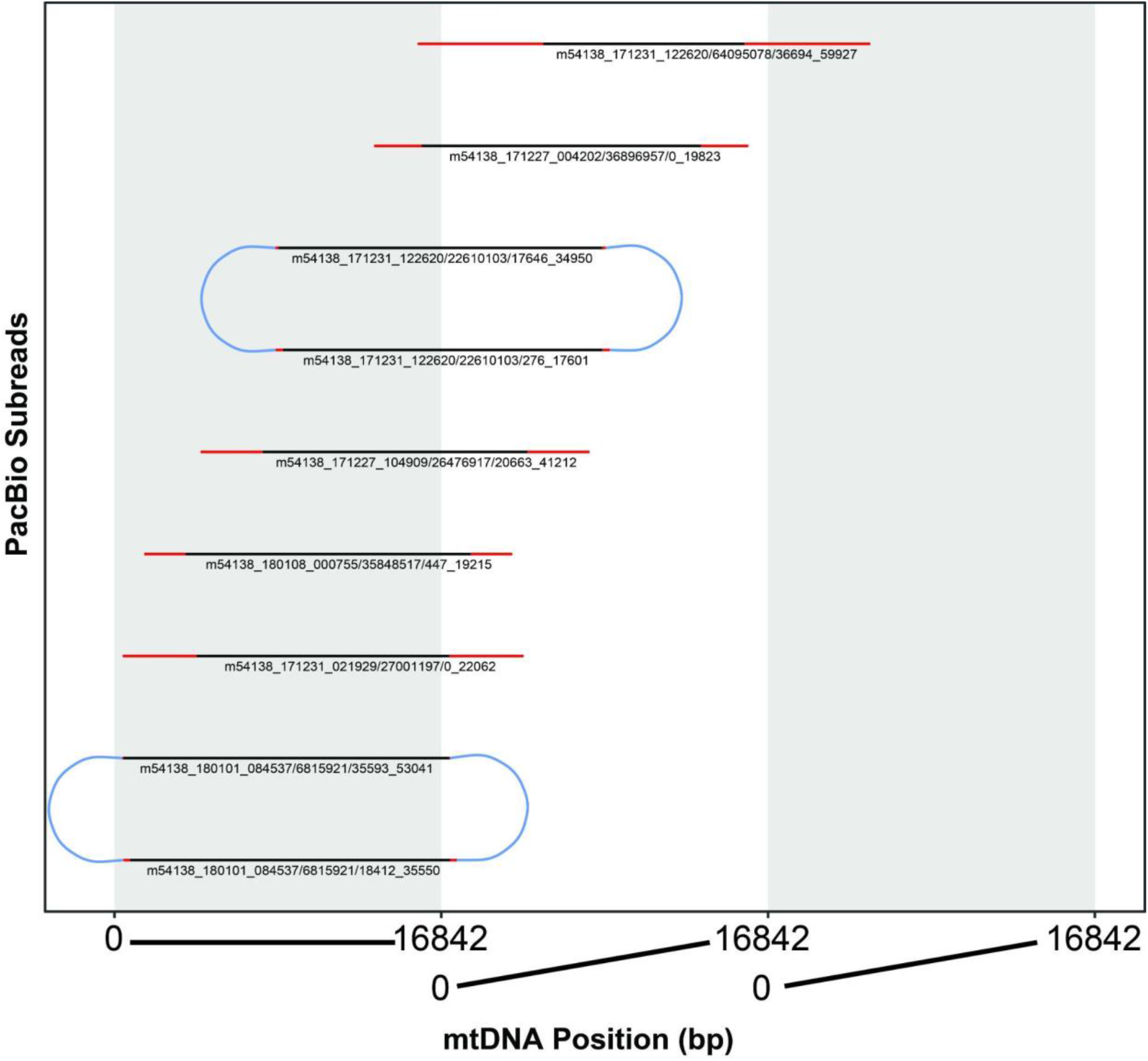
PacBio reads that captured more than a single mitochondrial genome in a single molecule. Black bars represent regions sequenced only once, and red regions of bars represent regions sequenced multiple times on the same molecule. Blue semi-circles connect consecutive subreads.<colcnt=1>

